# Universal brain signature of emerging reading in two contrasting languages

**DOI:** 10.1101/2019.12.18.881672

**Authors:** Katarzyna Chyl, Bartosz Kossowski, Shuai Wang, Agnieszka Dębska, Magdalena Łuniewska, Artur Marchewka, Marek Wypych, Mark van den Bunt, William Mencl, Kenneth Pugh, Katarzyna Jednoróg

## Abstract

Despite dissimilarities among scripts, a universal hallmark of literacy in adults is the convergent brain activity for print and speech. Little is known, however, how early it emerges. Here we compare speech and orthographic processing systems in two contrasting languages, Polish and English, in 100 7-year-old children performing identical fMRI tasks. Results show limited language variation, with speech-print convergence evident mostly in left fronto-temporal perisylvian regions. Correlational and intersect analyses revealed subtle differences in the strength of this coupling in several regions of interest. Specifically, speech-print convergence was higher for transparent Polish than opaque English in right temporal area, associated with phonological processing. Conversely, speech-print convergence was higher for English than Polish in left fusiform, associated with visual reading. We conclude that speech-print convergence is a universal marker of reading even at the beginning of reading acquisition while minor variations can be explained by the differences in the orthographic transparency.

## Introduction

Less than 6000 years ago writing systems began to develop to convey linguistic information through space and time. Despite striking dissimilarities among writing systems in regularity, frame and arrangement, they all represent the units of a spoken language. Irrespective of the writing system, reading depends on access to existing brain regions dedicated to the processing of spoken words. In consequence, the convergence of the speech and print processing systems onto a common neural network emerges as an invariant and universal signature of literacy proficiency (Rueckl et al., 2015). Whether the orthography is transparent or opaque, logographic or alphabetic - perisylvian regions in inferior frontal (IFG) and superior and middle temporal (STG/MTG) cortices were consistently co-activated by both spoken and written words in skilled adult readers of English, Spanish, Chinese and Hebrew (Rueckl et al., 2015). The authors argued that the invariance in speech-print convergence is the result of biological constraints imposed by perisylvian specialization for speech and natural language processing, and the need to use these specialized systems for print comprehension. Only subtle differences in the relative strength of speech-print convergence in several regions of interest were found between the languages. Particularly, speech-print convergence was slightly higher for transparent Spanish than opaque English and Hebrew in left supramarginal gyrus (SMG) and supplementary motor area, both associated with phonological processing (Herman et al., 2013). Conversely, speech-print convergence was higher for English and Hebrew relative to Spanish in several regions including left angular, fusiform (FG) and inferior temporal gyri (ITG) implicated in lexical-semantic processing in speech (Hickok & Poeppel, 2007) and in print (Pugh et al., 2010). Cross-lingual differences in speech-print convergence may be particularly pronounced at the initial stages of reading acquisition, where adequate orthography-phonology binding may be more beneficial in transparent languages.

To acquire reading, a child needs to master the ability to rapidly and accurately map letters to existing phonological representations (Wagner & Torgesen, 1987). Literacy acquisition reorganizes the brain (Dehaene et al., 2015), one example being the emergence of speech-print convergence (Chyl et al., 2018; Preston et al., 2016). While beginning readers of transparent Polish showed speech-print convergence in bilateral IFG and STG/MTG, it was absent in pre-readers matched for demographics (Chyl et al., 2018). Furthermore, in readers a positive correlation between convergence and reading skill was found in the left STG/MTG. Similarly, in English beginning readers reading readiness (as indexed by phonological awareness) was correlated with greater spatial speech-print convergence in the left STG/MTG (Frost et al., 2009). Importantly, the extent of the print-speech convergence can predict reading performance achieved one (Marks et al., 2019) or two years later (Preston et al., 2016) in English beginning readers. Regularity or orthographic transparency, a parameter that indicates how regular letter-phoneme correspondences are in the given script is a well-known factor influencing reading acquisition. Children learning to read in opaque orthographies are slower in acquiring this skill than children learning to read in transparent orthographies (Ziegler & Goswami, 2005) and thus might show lower spatial speech-print convergence. The orthographic depth hypothesis (Katz and Frost, 1992) as well as the psycholinguistic grain size theory (Ziegler and Goswami, 2005) suggest that learning to read based on phonological decoding is more advantageous for transparent orthographies and that whole word recognition is relatively more helpful for opaque scripts.

Using three complementary analytic approaches we examined print and speech processing networks and their convergence in 100 young users of two contrasting languages: opaque English and fairly transparent Polish (Schuppert, 2017), performing an identical fMRI language localizer. We expected that the general pattern of activity for print and speech will be similar across two languages, with speech-print convergence present in IFG and STG/MTG. Orthographic transparency effects should occur in regions related to phonological decoding with higher speech-print coupling in Polish than English, while the reversed pattern is expected in regions involved in visual word recognition.

## METHODS

### Participants

Inclusion criteria for the Polish sample were as follow: typical IQ, birth at term (>37 weeks), right-handedness, monolingualism, no history of neurological or language impairments and good quality of the fMRI scan (< 20% of motion-affected volumes identified with ART toolbox, see below for details). All English-speaking children who met the Polish inclusion criteria were included in the analysis (50 out of 82 collected datasets). Polish-speaking children were a part of the larger cohort (N = 120), and were matched pairwise with their American peers for age, word reading efficiency (N of words read correctly per minute) and a time gap between scan and behavioural test using the Hungarian optimization algorithm (cf. Chyl et al., 2018) to reduce group differences. As a result, data from 50 Polish (M age = 7.11, SD = 0.99, min = 5.41, max = 9.21) and 50 American (M age= 6.95, SD = 0.98, min = 4.75, max = 8.93) were selected for the current analysis. This sample size resulted in power higher than 80% for the fMRI analyses (Desmond & Glover, 2002). Similarly, this sample size gave us 80% power for detecting medium and large effects (Cohen’s d >= 0.50) in between-group comparisons, as revealed with G*Power (Faul, Erdfelder, Buchner, & Lang, 2009). All procedures were approved by the ethics committees in Poland (University of Warsaw Ethic Committee) and United States (Yale University School of Medicine). All parents gave written informed consent to the study and children agreed orally in compliance with human subjects protection and Helsinki Declaration guidelines.

### Behavioral measures

Word reading and pseudoword reading were tested with the Decoding Test (Polish; (Szczerbiński & Pelc-Pękała, 2013)) and Test of Word Reading Efficiency (English; (Torgesen, Wagner, & Rashotte, 2012)), and the raw score was scaled to the words per minute (WPM) measure. Since tests were not perfectly balanced for length, i.e. English words in the tests were shorter than Polish items, we estimated also letter per second measure. Rapid automatized naming (RAN) was tested with object naming subtest of Rapid Naming Test (Polish; (Fecenec, Jaworowska, Matczak, Stańczak, & Zalewska, 2013)) and Comprehensive Test of Phonological Processing (English; (R. K. Wagner, Torgesen, Rashotte, & Pearson, 2013)). Here, raw scores were scaled to the items per second score. In this subscale, all items in both languages were one-syllable words. On both sites the subscale of color naming was also applied, but since the Polish color names were longer than English (2.6 syllables on average versus 1.25), we did not include this measure in the analyses. Phonological awareness (PA) was examined with the phoneme deletion test (Polish; (Szczerbiński & Pelc-Pękała, 2013)) and (English; (R. K. Wagner et al., 2013)) and transformed into the normalized z-scores for each group. These PA tests had different instructions, items and timing so no direct comparison between languages was performed. Additionally, maternal and paternal education represented by the highest obtained grade (scaled to the 1-7 scale in both groups) was compared between the groups.

### fMRI and task procedure

Before the scanning session, children at both sites were familiarized with the task and scanner environment in a mock-scanner. Identical fMRI paradigm was used at both sites for print and speech activations localization (Malins et al., 2016). The event-related task consisted of four stimulus conditions: (1) printed real words, (2) spoken real words, (3) printed symbol strings, and (4) noise-vocoded spoken words to minimize phonetic content. Conditions (3) and (4) can be considered as low-level nonlinguistic control conditions that are matched in physical characteristics to the printed linguistic stimuli (length and visual complexity on screen) and to the spoken linguistic stimuli (dynamic frequency and amplitude content). However, linguistic content has been eliminated (orthographic and phonetic, respectively). This design activates the language network, and is sensitive to individual differences in reading skills in both adults (Malins et al., 2016) and children (Chyl et al., 2018). Polish children were asked to pay attention to the stimuli, but no explicit task was given to the participants. American children were also asked to pay attention to the stimuli and informed that after the task two simple recognition questions would be asked (e.g. „Did you hear the word „banana”?”). This step was introduced in order to make sure that children were focused on the task. However, reading should occur implicitly even without explicit instruction to read (Price, Wise, & Frackowiak, 1996) and listening is automatic as well.

On each trial, four different stimuli from the same condition were presented in rapid succession in a ‘tetrad’, designed to evoke strong activation within a relatively short imaging time. Each visual stimulus was presented for 250 ms, followed by a 200 ms blank screen, whereas each auditory stimulus was allowed 800 ms to play out. ‘Jittered’ intertrial intervals were employed with occasional ‘null’ trials resulting in ITIs ranging from 4 to 13 s (6.25 s on average). The task was performed in two runs, each lasting 5:02 minutes. All conditions were presented in each run, with 48 trials per run presented pseudorandomly, with restriction not to repeat one condition more than three times in a row. This resulted in 24 total trials per condition, and 96 total stimuli per condition. Stimuli were presented using Presentation software (Neurobehavioral Systems, Albany, CA) in Poland and E-Prime software in the United States.

### fMRI data acquisition

fMRI data at each site were acquired on Siemens 3T Magnetom Trio scanners using similar whole-brain echoplanar imaging sequences, 12-channel head coil (32 slices, slice-thickness 4 mm, TR = 2,000 ms, TE = 30 ms, FOV = 220×220 mm2, matrix size = 64 x 64, voxel size = 3 x 3 x 4). There was a difference in the flip angle parameter (Polish = 80°, American = 90°). Anatomical data was acquired using a T1 weighted MP-RAGE sequence (176 slices, slice-thickness = 1 mm, TR = 2,530 ms, TE = 3.32 ms, flip angle=7°, matrix size=256*256, voxel size= 1×1×1 mm). Generalized Autocalibrating Partial Parallel Acquisition (GRAPPA) acceleration was used at the Polish site (iPAT = 2), but not at the American site. To correct scanner differences, we performed iterative smoothness equalization and included signal-to-fluctuation-noise-ratio (SFNR) as a covariate in all between group comparisons (Friedman, Glover, & Fbirn Consortium, 2006).

### fMRI data processing and analysis

The preprocessing and analyses were performed using SPM12 (Wellcome Trust Center for Neuroimaging, London, UK) and AFNI version 17.3.09 (Cox, 1996). In SPM12, images were realigned to the first functional volume. Then structural images from single subjects were coregistered to their mean functional images. Coregistered anatomical images were segmented using pediatric tissue probability maps (generated with Template-O-Matic toolbox). Next, DARTEL was used to create a group-specific template and flow fields based on segmented tissues (Ashburner, 2007). Functional images were normalized to MNI space with 2×2×2mm voxel size using compositions of flow fields and a group-specific template. Next, in the univariate analyses, Gaussian spatial smoothing was performed using the 3dBlurtoFWHM option in AFNI, which allows for the „adaptive smoothing” method, and the data were smoothed to equalize estimated FWHM at 10 mm. The data were modeled using the canonical hemodynamic response function convolved with the experimental conditions and fixation periods. Movement regressors were added to the design matrix using ART toolbox to reject motion-affected volumes surpassing the movement threshold of 3 mm and a rotation threshold of 0.05 radians. On average 4.02 volumes were removed in the US, and 6.74 in PL samples, with non-significant difference between the groups.

To examine speech-print convergence we applied three different analytic approaches: intersect maps for print and speech on the whole brain and in selected regions of interest (ROIs), correlation analysis between brain activation to print and speech in selected ROIs and representational similarity analysis (*RSA*). Selection of ROIs was guided by the results on skilled adults (Rueckl et al., 2015) as well as meta-analyses of reading studies (Linkersdörfer, Lonnemann, Lindberg, Hasselhorn, & Fiebach, 2012; Richlan, 2012). Eight ROIs were included in the analyses: left and right STG/MTG, left and right IFG - with additional division to pars opercularis and pars triangularis in the left hemisphere (L IFG_oper and L IFG_tri, respectively), left SMG, left ITG and left FG. The ROIs were created using Anatomical Automatic Labeling (AAL) atlas (Tzourio-Mazoyer et al., 2002) masked with the functional activation defined as a sum of all activated regions for all contrasts of interest from both groups. Left angular gyrus and right SMG also reported by Rueckl and colleagues (Rueckl et al., 2015) were outside the activation mask and thus were not included as ROIs. In the ROI analyses we applied conservative Bonferroni correction for multiple comparisons to avoid false positives (i.e. p<0.05/8 = p≤0.00625).

Independent samples t-tests identified voxels that were significantly active at *P* < 0.005, FDR cluster corrected, for print and speech, print>symbols and speech>vocoded speech, separately for the two groups. Group conjunctions were explored based on conjunction null logic (Friston, Penny, & Glaser, 2005) in which we identified voxels that were significantly active at *P* < 0.005, FDR-corrected, for both PL and US in 4 conditions: print, speech, print>symbols and speech>vocoded speech.

To examine language differences within each anatomical ROI, we created a metric of speech-print convergence based on coactivation, defined as the total number of voxels for each participant that were significantly activated (p <0.05) both for speech and print (conjoint probability p < 0.0025; Frost et al., 2009; Marks et al., 2019; Preston et al., 2016). In addition, the number of voxels activated at p <0.05 across the functional mask defined as a sum of all activated regions for all contrast of interest from both groups for 1) spoken or 2) printed stimuli were computed to control for the relative degree of brain activation for each participant and together with 3) local SFNR were used as regressors of no interest.

In the correlation analysis, regression parameter estimates (averaged within the ROIs) for print and speech were used to compute r-Pearson correlation coefficients across subjects in each group. Correlation coefficients were then compared between languages using the Fisher r-to-z transformation.

The searchlight RSA was conducted for each subject by using RSA toolbox (Kriegeskorte, Mur, & Bandettini, 2008), and was constrained in gray matter with a gray matter mask generated from AAL template (Tzourio-Mazoyer et al., 2002). After obtaining trial-wise estimates with beta-series regression (Rissman, Gazzaley, & D’Esposito, 2004), 96 trial-wise beta images were used to assess representational dissimilarity between every pair of trials within a spherical searchlight kernel with 9 mm radius centered at each gray matter voxel, resulting in a representational dissimilarity matrix (RDM) map in which each voxel contains a 96 by 96 RDM. Specifically, the representational dissimilarity between a pair of trials was estimated by one minus Pearson correlation (1 - r) where the correlation was calculated between beta values within a searchlight kernel. The speech-print convergence model was constructed as a RDM where the printed and spoken words are regarded as identical so that the trial pairs of real words hold highest similarity (valued 0 in RDM) while other trial pairs yield lowest similarity (valued 1 in RDM). The representational similarity between neural representation and the speech-print convergence model were estimated by calculating Spearman’s rho between the RDM maps and the model RDM for each voxel. The resulting Spearman’s rho maps were then Fisher-z transformed and submitted to second-level statistical tests. All RSA results are presented on the voxel threshold p < 0.005, FDR cluster corrected.

Additionally, activation to print only or speech only, as well as print>symbols and speech>vocoded speech was compared between the languages within the selected ROIs, corrected for SFNR. Whole-brain group comparisons were not performed, as they are potentially more susceptible to cross-scanner differences, and could result in differences in regions outside the canonical reading and speech networks (Rueckl et al., 2015).

Behavioural data, ROI data, parameters of the items used in fMRI experiment as well as the experimental protocols used at both sites are available online (https://osf.io/982ks).

Figures were prepared with Nipype (Gorgolewski et al., 2011).

## RESULTS

### Behavioral results

Demographics and test performance is presented in Table 1. Since the groups were matched for reading, no differences were found for word reading score. However, independent samples t-test showed significant differences between Polish and American children in the estimated scores of letters in pseudowords read per second, with Polish children reading more efficiently than American. Since no difference was found in the pseudowords per minute, this result reflects the differences in test items, as pseudowords used in US group were shorter. There was no difference between the fathers’ education, but mothers of the PL group obtained higher level of education.

**Table 1.** Demographics and test performance in Polish (PL) and American (US) children: Means, (Standard Deviations) and [95% CIs].

|  |  | PL (n=50) | US (n=50) | test | p-value | Cohen's d |
| --- | --- | --- | --- | --- | --- | --- |
| Demographic measures | Age in years | 7.11 (0.98) [6.84; 7.38] | 6.95 (0.99) [6.68; 7.22] | t = 0.816 | ns | 0.16 |
|  | Sex: M = males, F = females | 18 M, 32 F | 22 M, 28 F | Chi<sup>2</sup> = 0.667 | ns |  |
|  | Mother Education | 6.46 (0.96) [6.19; 6.73] | 5.78 (1.09) [5.48; 6.08] | t = 3.332 | p = 0.001 | 0.66 |
|  | Father Education | 6.00 (1.41) [5.61; 6.39] | 6.02 (1.17) [5.70; 6.34] | t = 0.076 | ns | 0.02 |
| Behavioral measures | Word reading: words read correctly per minute | 41.96(30.81) [33.42; 50.50] | 50.58 (33.81) [41.21; 59.95] | t = 1.332 | ns | 0.27 |
|  | Pseudoword reading: pseudowords read correctly per minute | 27.02 (16.70) [22.39; 31.65] | 23.38 (20.01) [17.83; 28.93] | t = 0.987 | ns | 0.20 |
|  | Letters in words read per second (estimated) | 3.28 (2.90) [2.48; 4.08] | 3.33 (2.78) [2.56; 4.10] | t = 0.086 | ns | 0.02 |
|  | Letters in pseudowords read per second (estimated) | 2.35 (1.78) [1.86; 2.84] | 1.31 (1.41) [0.92; 1.70] | t = 3.266 | p = 0.002 | 0.65 |
|  | RAN: objects named per second | 0.86 (0.20) [0.80; 0.92] | 0.95 (0.23) [0.89; 1.01] | t = 2.474 | p = 0.015 | 0.42 |

ns - not significant

### fMRI results

#### Language-independent activation

Figure 1 and Table 2 reports the results of the group conjunction analysis revealing language-independent networks for printed and spoken word recognition. For print, the regions that were commonly employed by Polish and American children were bilateral occipital, frontal and temporal cortex. Print specific (print > symbols) activation common for both groups was present solely in the left IFG and precentral gyrus (PrCG). For speech and speech specific (speech > vocoded) conditions both groups activated bilateral temporal and frontal cortex, but speech specific activation was less extensive.

**Table 2.**
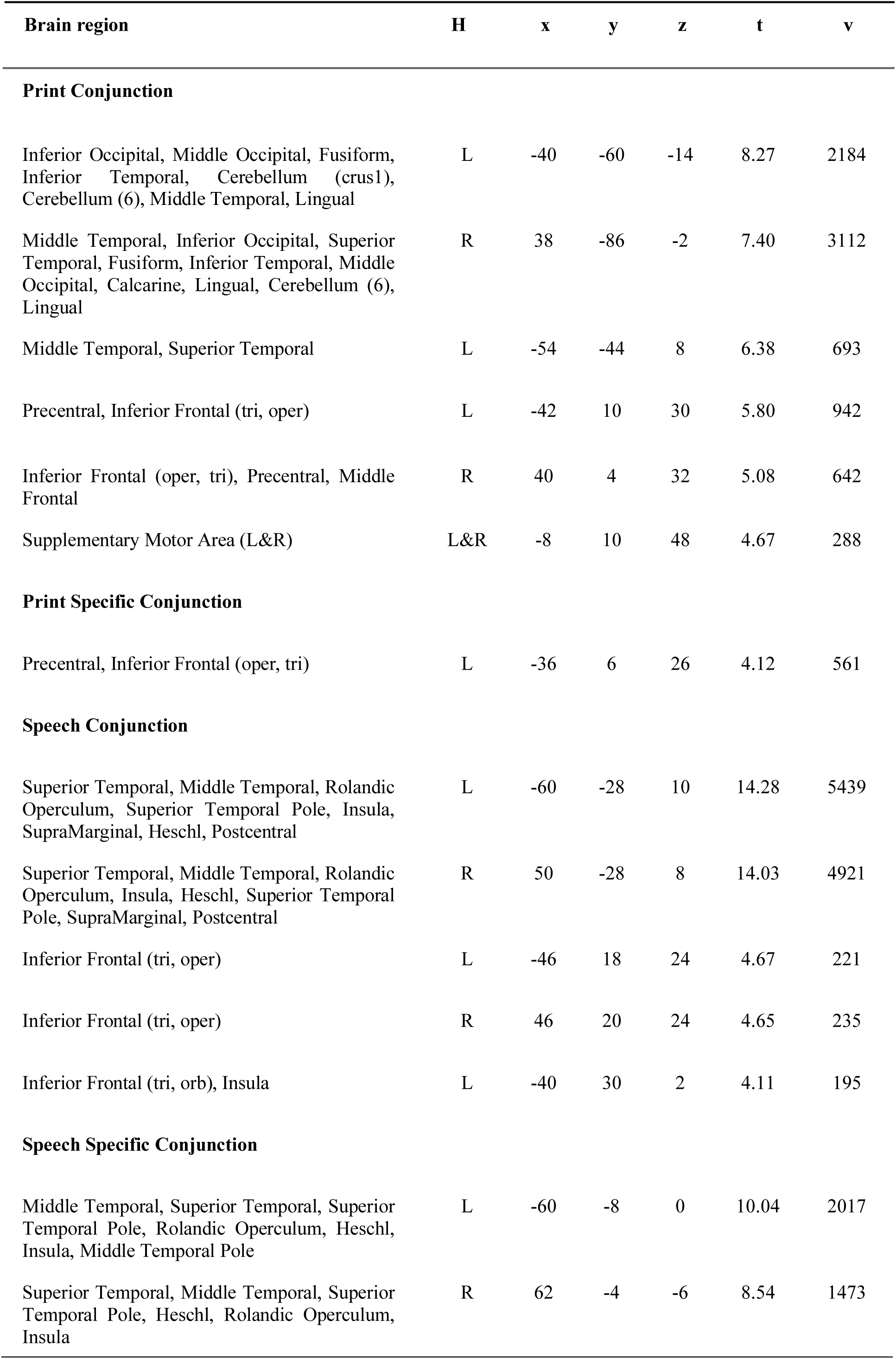
Brain regions that are active in both groups for Print, Speech, Print Specific (Print > Symbols) and Speech Specific (Speech > Vocoded). Hemisphere (H), coordinates (x, y, z), t statistic for the peak (t) and number of voxels (v) is reported.

| Brain region | H | x | y | z | t | v |
| --- | --- | --- | --- | --- | --- | --- |
| **Print Conjunction** |  |  |  |  |  |  |
| Inferior Occipital, Middle Occipital, Fusiform, Inferior Temporal, Cerebellum (crus1), Cerebellum (6), Middle Temporal, Lingual | L | -40 | -60 | -14 | 8.27 | 2184 |
| Middle Temporal, Inferior Occipital, Superior Temporal, Fusiform, Inferior Temporal, Middle Occipital, Calcarine, Lingual, Cerebellum (6), Lingual | R | 38 | -86 | -2 | 7.40 | 3112 |
| Middle Temporal, Superior Temporal | L | -54 | -44 | 8 | 6.38 | 693 |
| Precentral, Inferior Frontal (tri, oper) | L | -42 | 10 | 30 | 5.80 | 942 |
| Inferior Frontal (oper, tri), Precentral, Middle Frontal | R | 40 | 4 | 32 | 5.08 | 642 |
| Supplementary Motor Area (L&R) | L&R | -8 | 10 | 48 | 4.67 | 288 |
| **Print Specific Conjunction** |  |  |  |  |  |  |
| Precentral, Inferior Frontal (oper, tri) | L | -36 | 6 | 26 | 4.12 | 561 |
| **Speech Conjunction** |  |  |  |  |  |  |
| Superior Temporal, Middle Temporal, Rolandic Operculum, Superior Temporal Pole, Insula, SupraMarginal, Heschl, Postcentral | L | -60 | -28 | 10 | 14.28 | 5439 |
| Superior Temporal, Middle Temporal, Rolandic Operculum, Insula, Heschl, Superior Temporal Pole, SupraMarginal, Postcentral | R | 50 | -28 | 8 | 14.03 | 4921 |
| Inferior Frontal (tri, oper) | L | -46 | 18 | 24 | 4.67 | 221 |
| Inferior Frontal (tri, oper) | R | 46 | 20 | 24 | 4.65 | 235 |
| Inferior Frontal (tri, orb), Insula | L | -40 | 30 | 2 | 4.11 | 195 |
| **Speech Specific Conjunction** |  |  |  |  |  |  |
| Middle Temporal, Superior Temporal, Superior Temporal Pole, Rolandic Operculum, Heschl, Insula, Middle Temporal Pole | L | -60 | -8 | 0 | 10.04 | 2017 |
| Superior Temporal, Middle Temporal, Superior Temporal Pole, Heschl, Rolandic Operculum, Insula | R | 62 | -4 | -6 | 8.54 | 1473 |

**Figure 1.**
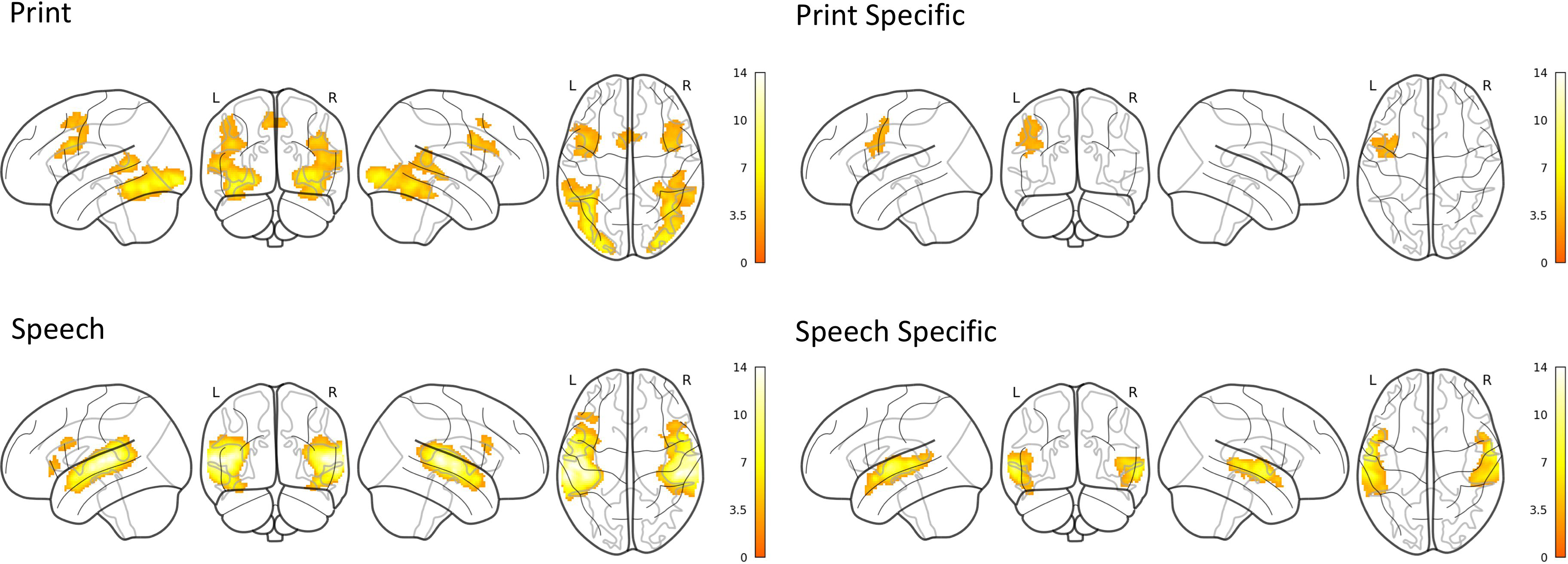
Group conjunctions showing brain regions that are active for both groups in Print, Print Specific (print > symbols), Speech, Speech Specific (speech > vocaded) (threshold for each contrast p < 0.005, FDR cluster corrected) for each language.

#### Speech-print convergence

Figure 2 presents regions active for print and speech (for details see Table S2), as well as regions convergently active for print and speech in both groups (Table 3). Whole brain convergence analysis for speech and print revealed activation in bilateral IFG and MTG/STG for both Polish and English with additional cluster of overlap in the right parietal cortex for American children. Speech and print specific intersection was visible only in Polish children in bilateral MTG/STG at the given threshold.

**Table 3.**
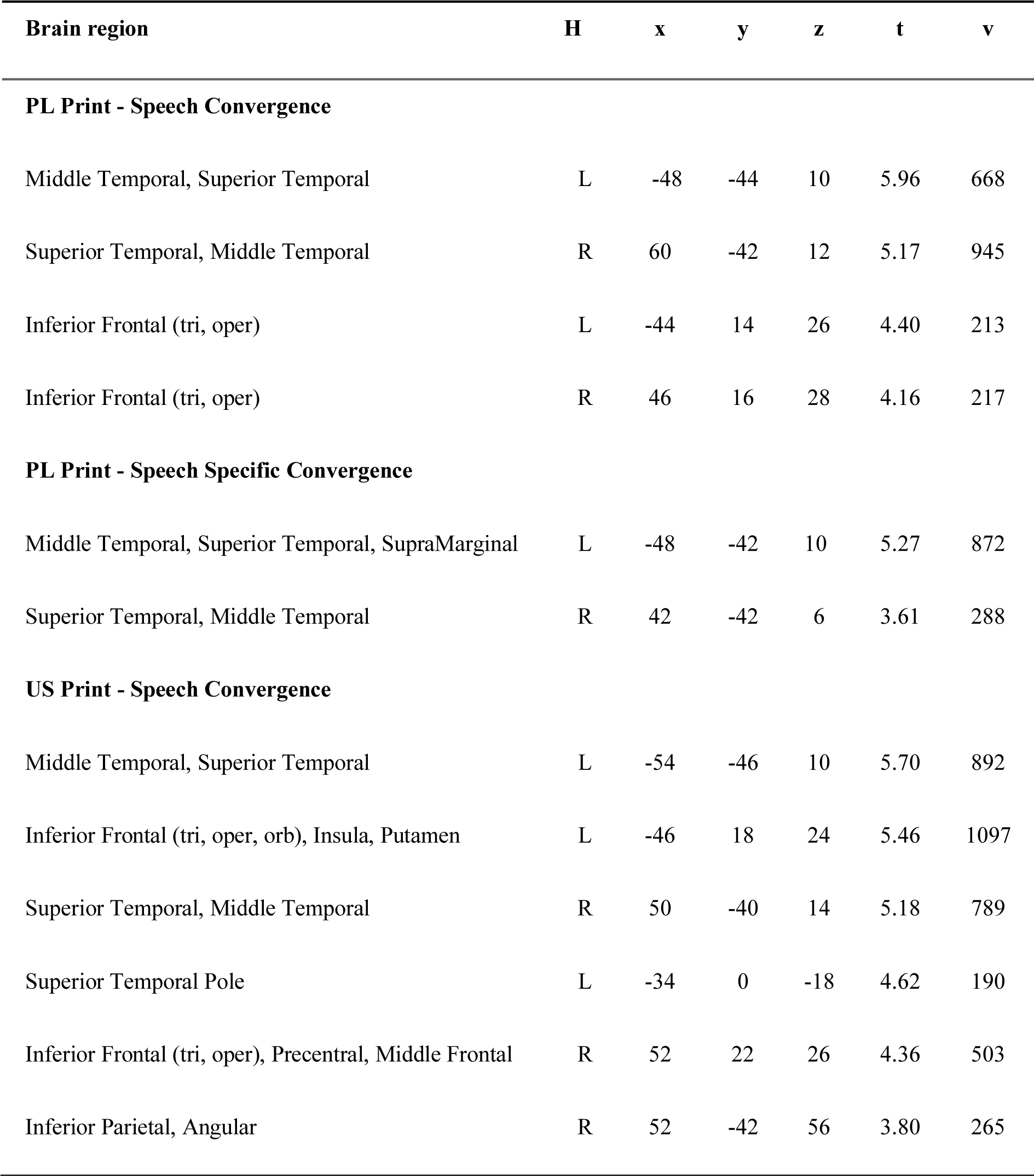
Print-Speech Convergence in Polish and American groups. Threshold for each contrast p < 0.005, FDR-corrected. Hemisphere (H), coordinates (x, y, z), t statistic for the peak (t) and number of voxels (v) are reported.

| Brain region | H | x | y | z | t | v |
| --- | --- | --- | --- | --- | --- | --- |
| **PL Print - Speech Convergence** |  |  |  |  |  |  |
| Middle Temporal, Superior Temporal | L | -48 | -44 | 10 | 5.96 | 668 |
| Superior Temporal, Middle Temporal | R | 60 | -42 | 12 | 5.17 | 945 |
| Inferior Frontal (tri, oper) | L | -44 | 14 | 26 | 4.40 | 213 |
| Inferior Frontal (tri, oper) | R | 46 | 16 | 28 | 4.16 | 217 |
| **PL Print - Speech Specific Convergence** |  |  |  |  |  |  |
| Middle Temporal, Superior Temporal, SupraMarginal | L | -48 | -42 | 10 | 5.27 | 872 |
| Superior Temporal, Middle Temporal | R | 42 | -42 | 6 | 3.61 | 288 |
| **US Print - Speech Convergence** |  |  |  |  |  |  |
| Middle Temporal, Superior Temporal | L | -54 | -46 | 10 | 5.70 | 892 |
| Inferior Frontal (tri, oper, orb), Insula, Putamen | L | -46 | 18 | 24 | 5.46 | 1097 |
| Superior Temporal, Middle Temporal | R | 50 | -40 | 14 | 5.18 | 789 |
| Superior Temporal Pole | L | -34 | 0 | -18 | 4.62 | 190 |
| Inferior Frontal (tri, oper), Precentral, Middle Frontal | R | 52 | 22 | 26 | 4.36 | 503 |
| Inferior Parietal, Angular | R | 52 | -42 | 56 | 3.80 | 265 |

**Figure 2.**
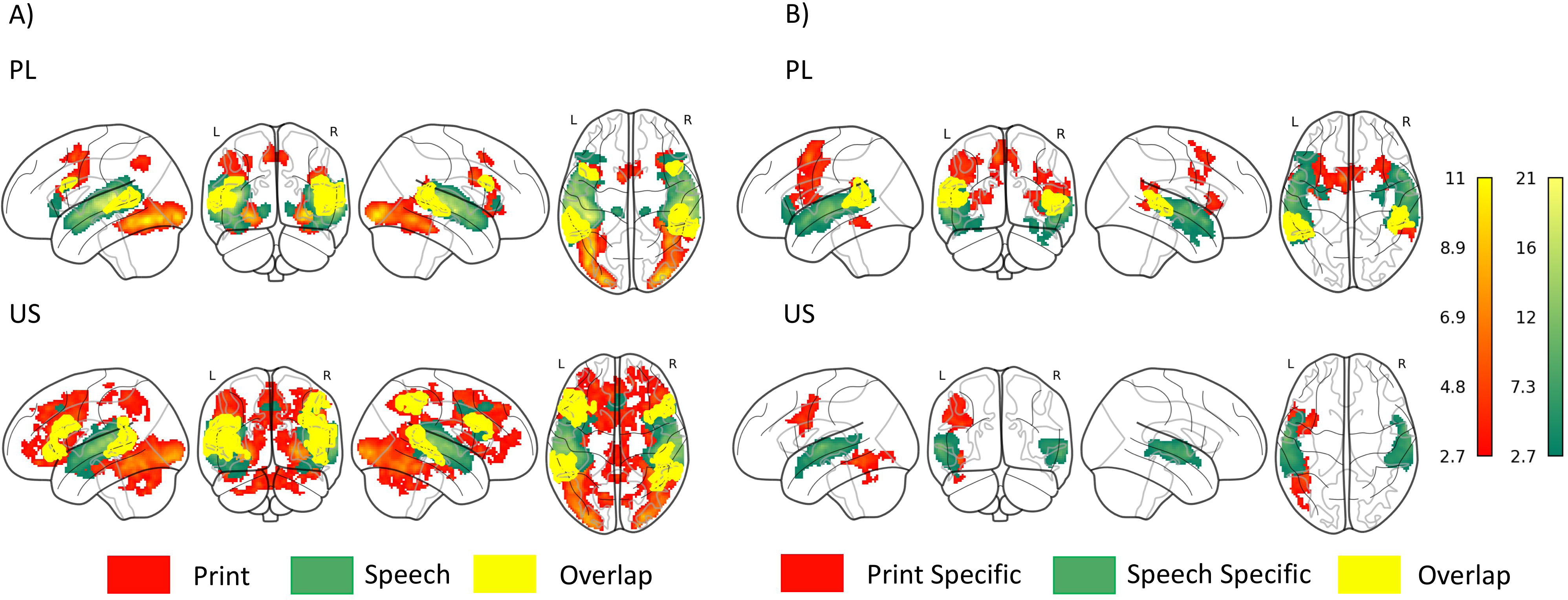
A) Intersect maps showing brain regions that are active for Print only (red), Speech only (green), or both (yellow) (threshold for each modality p < 0.005, FDR cluster corrected) for Polish (PL) and English (US). B) Intersect maps showing brain regions that are active for Print Specific (red), Speech Specific (green), or both (yellow) (threshold for each modality p < 0.005, FDR cluster corrected) for each language.

Individual convergence analysis within the ROIs revealed that speech-print convergence was higher for Polish than English in right STG/MTG (t(98) = 3.065, p = 0.003), while the reversed pattern was present in the left FG (t(98) = 2.979, p=0.004). No significant differences between the groups were found for speech or print specific convergences.

Similar results were observed in the brain activation correlation analysis within the ROIs (Figure 3). While the correlation between regression parameter estimates for print processing and speech processing in the left FG was significant in American children (r = 0.518 [0.282; 0.696], p<0.001) it did not reach significance in Polish children (r=0.259 [0; 0.501], p=0.07), however the difference between correlation coefficients was not significant (z=1.5; p=0.13). In case of the right STG/MTG, the correlation was significant in both languages (r=0.636 [0.438; 0.778], p<0.001 and r=0.301 [0.030; 0.537], p=0.034 for PL and US respectively), but was significantly higher in Polish than English (z=2.14; p=0.03). Additionally, the significant difference in the correlation coefficients was found in the left IFG (pars opercularis; z = 2.2, p = 0.028), with significant correlation found in PL (r = 0.626 [0.422; 0.770], p < 0.001) and at a trend level in US (r = 0.274 [0.00; 0.515], p = 0.054). Again, no significant correlations (surviving correction for multiple comparisons) were revealed for print and speech specific contrasts.

**Figure 3.**
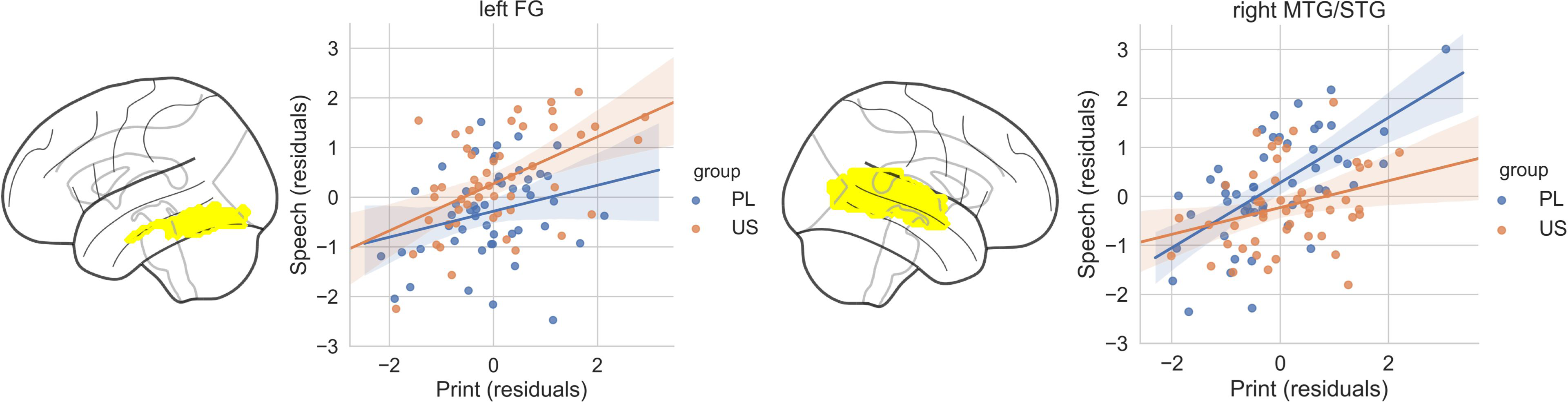
Scatter plots of the correlation between print and speech activation in representative areas showing greater convergence in right STG/MTG for more transparent Polish (Left), and in left fusiform gyrus (FG) for opaque English (Right). Fisher’s R-to-Z transform was performed to check the difference between the languages.

A high degree of similarity in speech-print convergence between Polish and American children was also revealed in RSA analysis (Figure 4 and Table 4). Again, the convergence as measured by similarity between brain response to speech and print was present in bilateral temporal regions and left frontal areas. No significant differences between the groups were found in RSA ROI analyses.

**Table 4.**
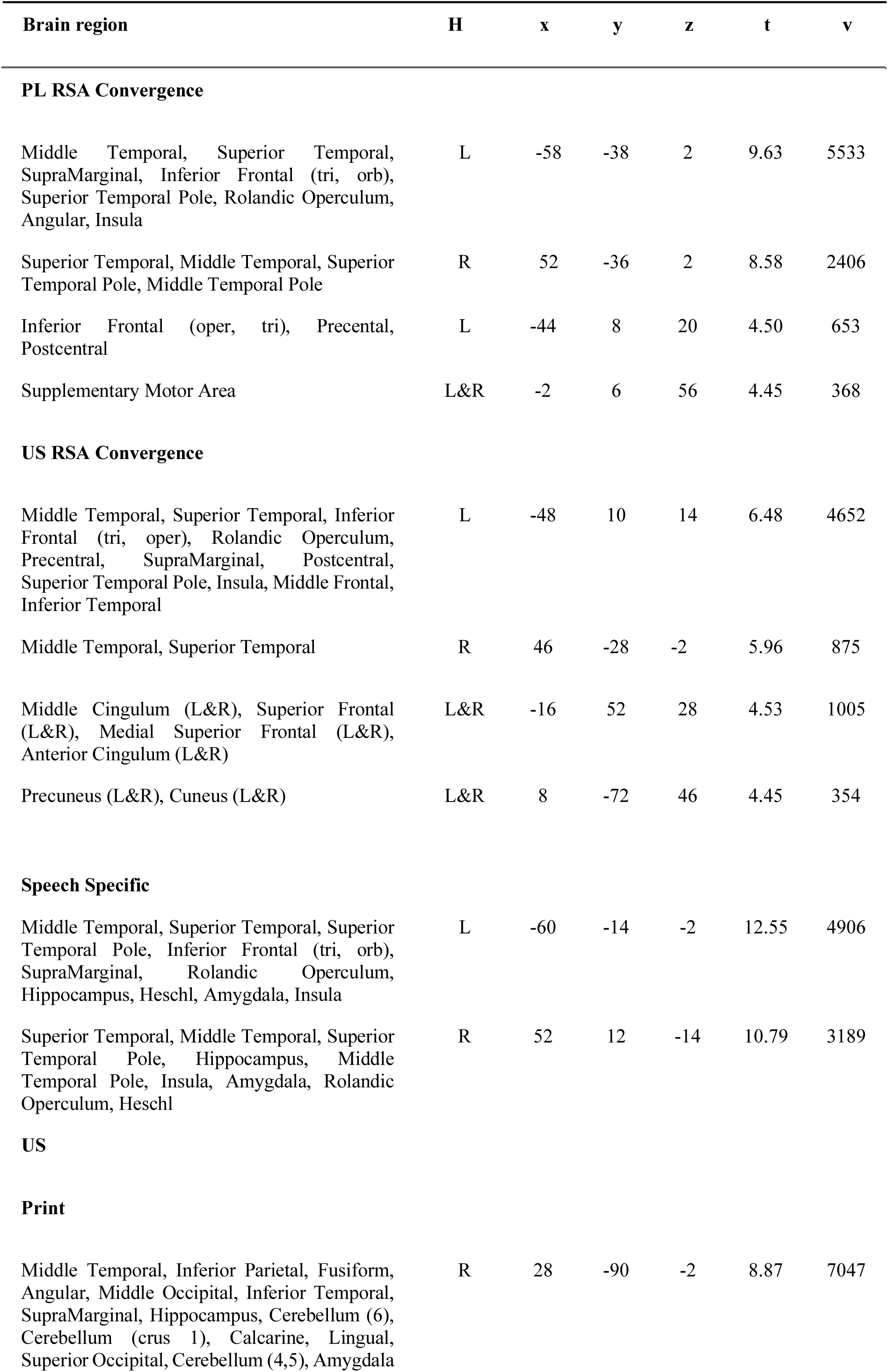
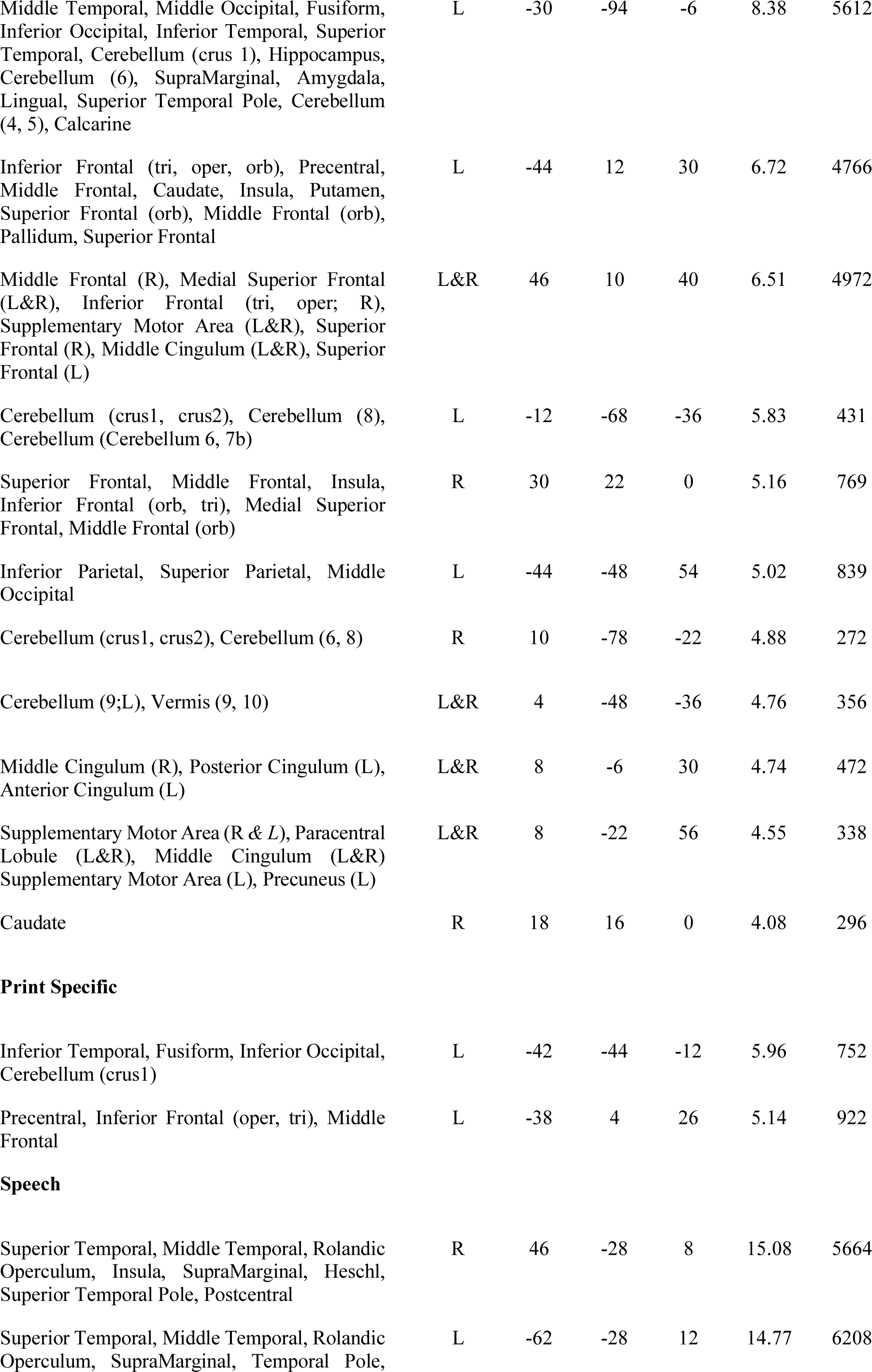
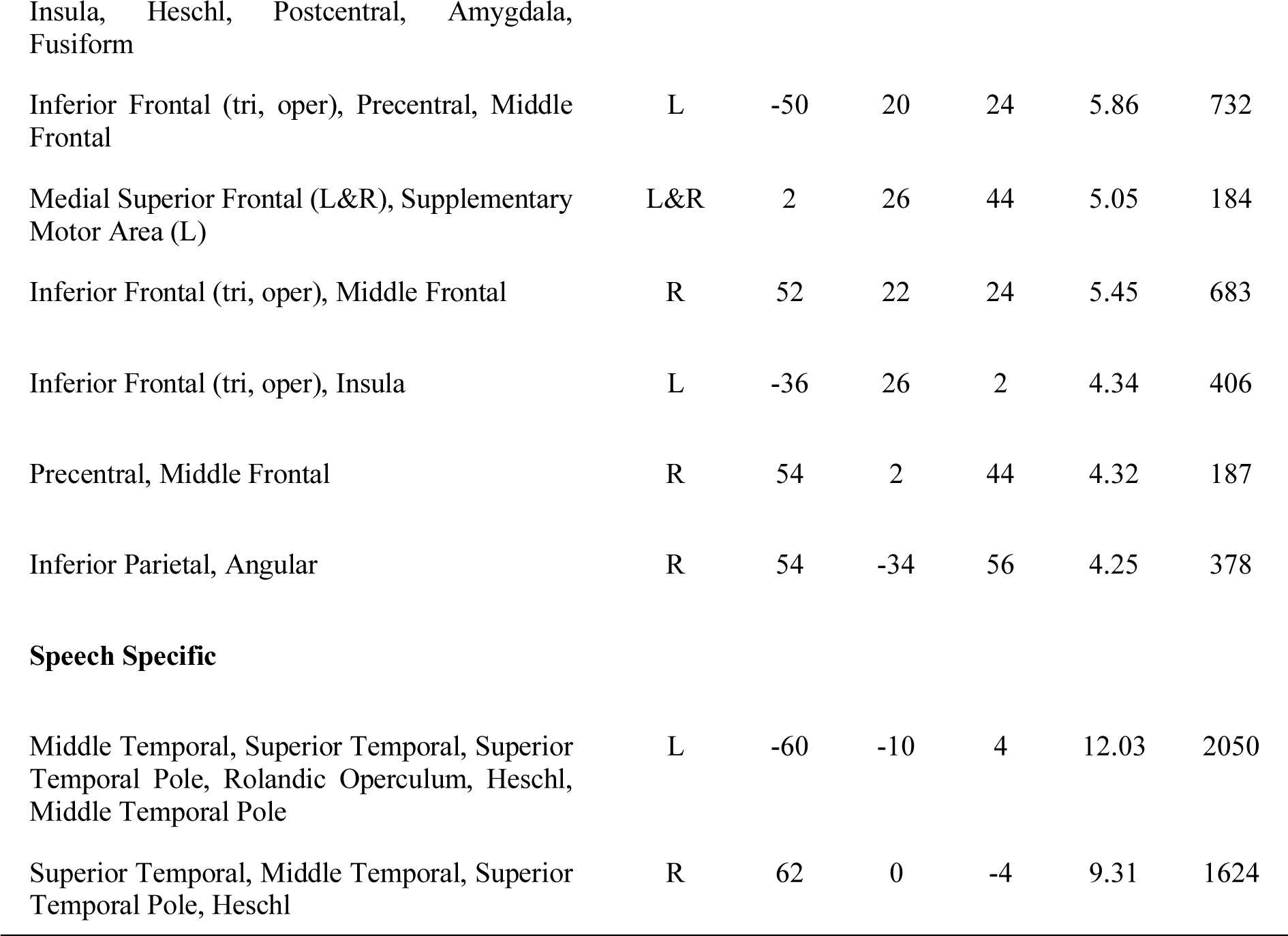
RSA Convergence maps in Polish and American groups. Threshold for each contrast p < 0.005, FDR-corrected. Hemisphere (H), coordinates (x, y, z), t-statistic for the peak (t) and number of voxels (v) are reported.

**Figure 4.**
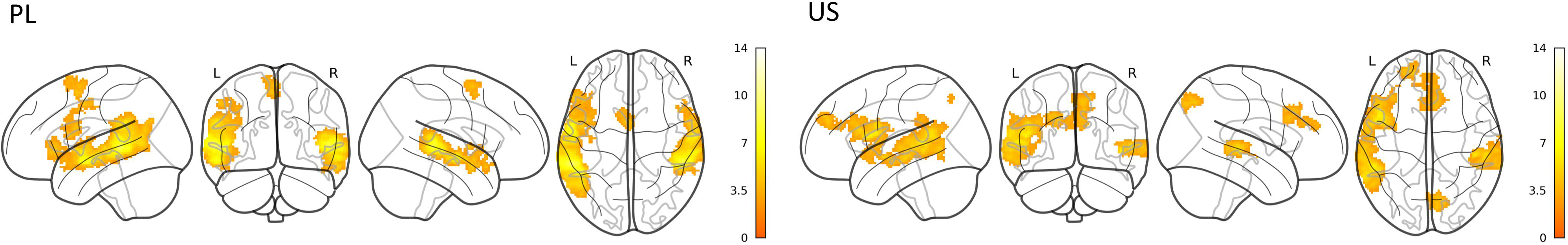
RSA convergence maps in Polish and American children (threshold p < 0.005, FDR cluster corrected).

#### Language-specific activation

Next, we examined group differences in activation to print only or speech only, as well as print>symbols and speech>vocoded speech within the selected ROIs. For visual conditions, only one significant difference was found, with English involving left IFG pars triangularis more than Polish in response to print (t(98) = 3.163, p < 0.002). In print specific condition no differences were found. For speech, English had higher activation than Polish in the left FG (t(98) = 3.167, p = 0.002) and ITG (t(98) = 4.243, p < 0.001), while left MTG/STG was more involved in Polish than English (t(98) = 3.280, p = 0.001). Polish produced also higher response in the left MTG/STG than English in speech specific condition (t(98) = 3.314, p = 0.001).

## Discussion

Here, we present how young beginning readers of Polish and English process spoken and printed words. We particularly focused on the aspect of conjoint processing of print and speech, a hallmark of the successful literacy acquisition (Chyl et al., 2018) and common for different languages in skilled adult readers (Rueckl et al., 2015). We also tested language-related similarities and differences in processing print and speech separately.

Our results show striking resemblance to previous findings (Rueckl et al., 2015), and demonstrate that incorporating print into the existing speech network is similar in contrasting languages, not only in adulthood but also at the beginning of reading acquisition. Bilateral IFG and MTG/STG were activated by print and speech in both Polish and American children. Complementary RSA analysis confirmed language invariant speech-print coactivation in the left IFG and bilateral MTG/STG. Speech-print convergence in the previous study (Rueckl et al., 2015) was additionally present in left parietal cortex, which may be related to the task demands. Here, we measured implicit activation to speech and print, while in previous study participants made semantic judgments. Nevertheless, we provide evidence that the core speech-print convergence is independent of reading experience and the fMRI task, at least for typical reading development.

When we tested the size of speech-print convergence in several ROIs of the language network, we found that Polish children had more convergent voxels in the right STG/MTG than American, while a reversed pattern was present in the left FG. These results were supported by the additional correlational analysis showing stronger speech-print correlations of neuronal activity in the right STG/MTG in Polish than English. In the left FG, the speech-print correlation was significant only in English, but not in Polish (though the difference between languages did not reach significance). Since STG/MTG is generally associated with phonological processing and left FG with lexical processing, our results support the predictions from both orthographic depth hypothesis (Frost and Katz, 1992) and the psycholinguistic grain size theory (Ziegler & Goswami, 2005). Polish children rely more on right STG/MTG using phonological decoding for reading, while American children reading in English rely more on whole word recognition. These findings are also in line with Rueckl et al. (2015) who found stronger print-speech coupling in the regions related to phonological processing - left SMG and SMA (Stoeckel et al., 2009) in orthographically transparent Spanish than in English and Hebrew. Orthographically opaque English and Hebrew had stronger convergence not only in left FG, but also in the left angular gyrus, MTG, ITG and IFG (pars triangularis), related to the semantic processing. In contrast to current findings, right STG and SMG also showed stronger correlations for the comparison of opaque versus transparent orthographies. Besides the potential influence of reading experience and employed task, some of the examined adult participants were multilingual (in contrast to currently examined monolingual children), which might have affected the pattern of brain activation. Nevertheless, the reported differences in speech-print convergence between beginning and skilled readers of contrasting orthographies are rather subtle, supporting the claim that reading network is deeply constrained by the organization of the brain network also at the beginning of reading acquisition.

Print stimulation in both languages evoked activity in bilateral inferior occipital, temporal and frontal areas, thus the classical network for reading (Dehaene et al., 2010; Martin et al., 2015). At the same time, print specificity (print>symbols) was found only in left IFG and PrCG in both groups. Engagement of the left IFG/PrCG in early reading was shown in both typical and struggling readers across different languages (Pollack, Luk, & Christodoulou, 2015) and was associated with phonological recoding (Pugh et al., 2010) or top-down cognitive control (Pollack et al., 2015). We previously showed that the left IFG/PrCG shows stronger activation to words in readers compared to age-matched pre-readers (Chyl et al., 2018) and its significance for reading increases with time and reading instruction (Chyl et al., 2019). Currently we demonstrate that PrCG/IFG activity is the only common word specific activation in young readers of two languages. Study on young German readers found that print>symbols induced activity in IFG and MTG (Bach et al., 2013); similar pattern was found in Polish, however the American group activated merely left hemisphere. We speculate that this result may be related to the similar orthographic transparency of Polish and German. However, in print>symbols comparison no significant differences between the groups were found. Only for print itself stronger involvement of the left IFG (pars triangularis) was found for English than Polish. This structure is often implicated in semantic processes of reading and stronger activation in the American cohort may reflect a stronger reliance on lexical-semantic processes.

Common speech activation was found in the bilateral temporal and frontal regions, while speech specific activation was limited to the bilateral temporal cortex. Similarly, Rueckl et al., (2015) examining adults showed that STG was active for speech regardless of language. It is not surprising, considering the biological constraints imposed by perisylvian specialization for speech. However, reading training was shown to reorganize these areas and enhance speech processing in planum temporale/STG (Monzalvo & Dehaene-Lambertz, 2013), and speech specific activity in the left STG was shown to correlate with reading efficiency in beginning readers (Chyl et al., 2018). Here, we found that Polish children engaged left STG/MTG stronger than American for both speech and speech specific contrasts. This result suggests that reorganization of the speech network is a consequence of reading acquisition proceeding faster and more easily in readers of a transparent script. An alternative explanation relates to the fMRI task material, as Polish words matched for frequency and length to American words had higher number of syllables and phonemes (Syllables: mean PL=1.28, mean US=1; t(382)=6.912, p<0.001; Phonemes: mean PL=3.85, mean US=3.54; t(382)=3.220, p=0.002) and it has been shown before that STG is particularly sensitive to these linguistic properties (Perrachione et al., 2017). Higher activation for American than Polish was found in the left FG and ITG, but only for speech. Activity of the ITG in response to speech was observed in 9-year olds but not pre-reading 6-year olds in the previous study (Monzalvo & Dehaene-Lambertz, 2013) and was explained as the sign of the orthographic influences on speech perception.

Current findings come from a multicenter study, and certain differences in both behavioral tests and fMRI data acquisition have to be acknowledged. We have tried to diminish potential sources of unwanted variance by carefully matching the subjects for demographics and reading skills and following FBIRN recommendations for handling multicenter fMRI data (Glover et al., 2012). However, we cannot exclude the possibility that not all of the confounding factors have been cancelled out.

In summary, we have demonstrated that in the two groups of children speaking different languages the neural pattern of print and speech processing is remarkably similar. Importantly, the speech-print convergence is present in both groups, yet again suggesting that incorporating orthographic processing into the speech pathways shaped by evolution is universal for different languages and scripts. However, orthographic transparency of the language may evoke different strategies in early reading, as suggested by the orthographic depth hypothesis (Katz & Frost, 1992). In our study American children showed stronger involvement of the fusiform gyrus for print and its stronger print-speech coupling, while the Polish children showed higher speech-print convergence in the right middle and superior temporal gyri, associated with phonological processing.

## Author Contributions

K. Jednoróg and K. Pugh developed the study concept. Together with W. Mencl they designed the experiment. K. Chyl together with A. Dębska and M. Łuniewska collected the PL data. Working under the supervision of K. Jednoróg, K. Chyl matched the groups and analyzed the data. K. Jednoróg and K. Chyl interpreted the data and drafted the manuscript. S. Wang, B. Kossowski and M. Wypych helped with data analysis. S. Wang drafted the methods section regarding RSA. A. Marchewka and M.R. van den Bunt helped with the interpretation of the results. All authors provided critical revisions and approved the final version of the manuscript for the submission.

## Acknowledgements

This work was funded by grants from the Polish Ministry of Science and Higher Education (IP2011 020271), the National Science Center (2014/N/HS6/03515, 2011/03/D/HS6/05584, 2014/14/A/HS6/00294), Eunice Kennedy Shriver National Institute of Child Health and Human Development (P01 HD 001994, P01 HD 070837) and National Institutes of Health (5R01HD086168-04, 5R37HD090153-03). The project was realized with the aid of CePT research infrastructure purchased with funds from the European Regional Development Fund as part of the Innovative Economy Operational Programme, 2007-2013. Funding sources were not involved in the experiment realization, data collection, data analysis, or writing of the report. The authors would like to thank all the families which participated in this study.

## Supplementary Materials

**Table S1.**
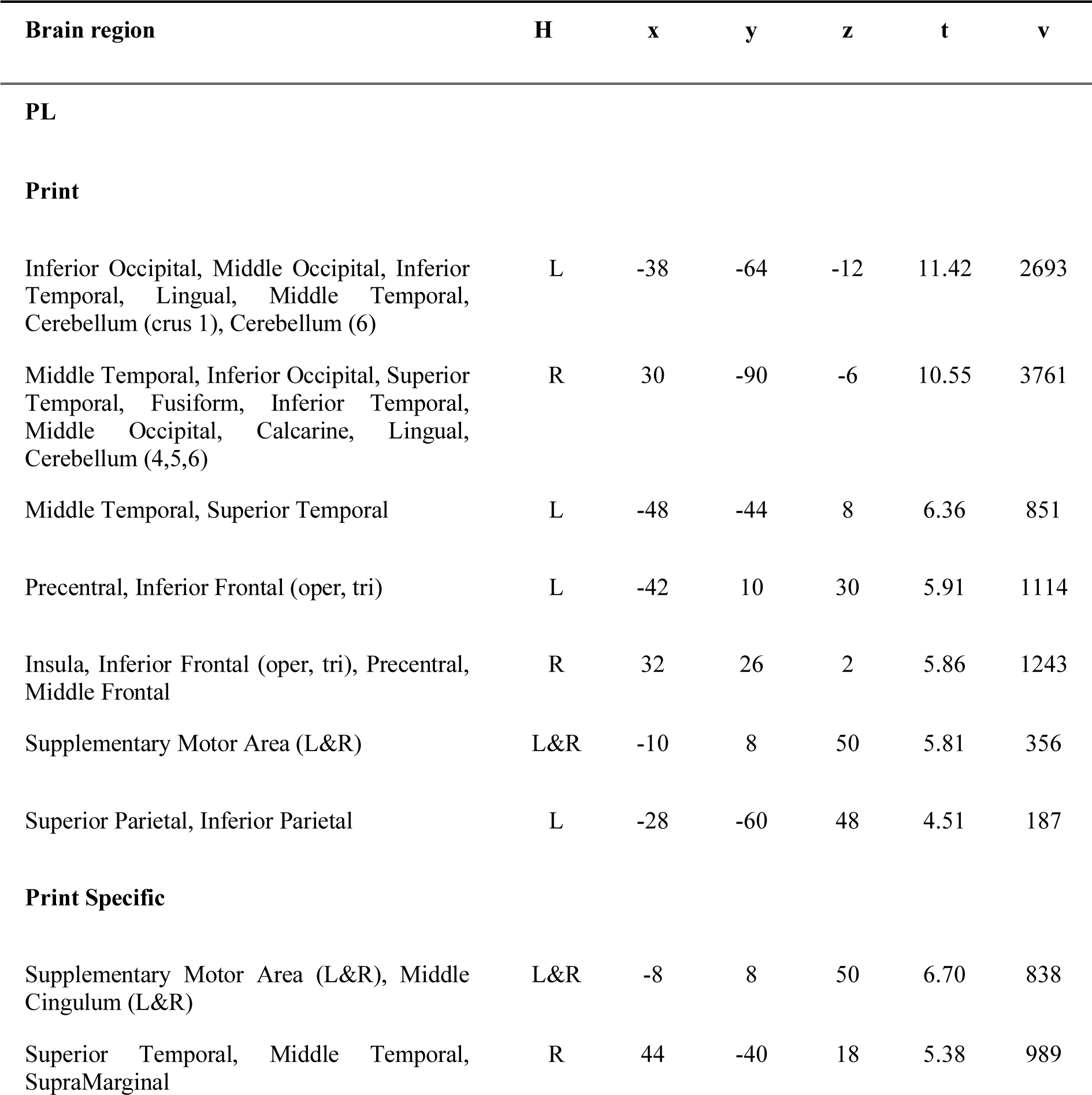
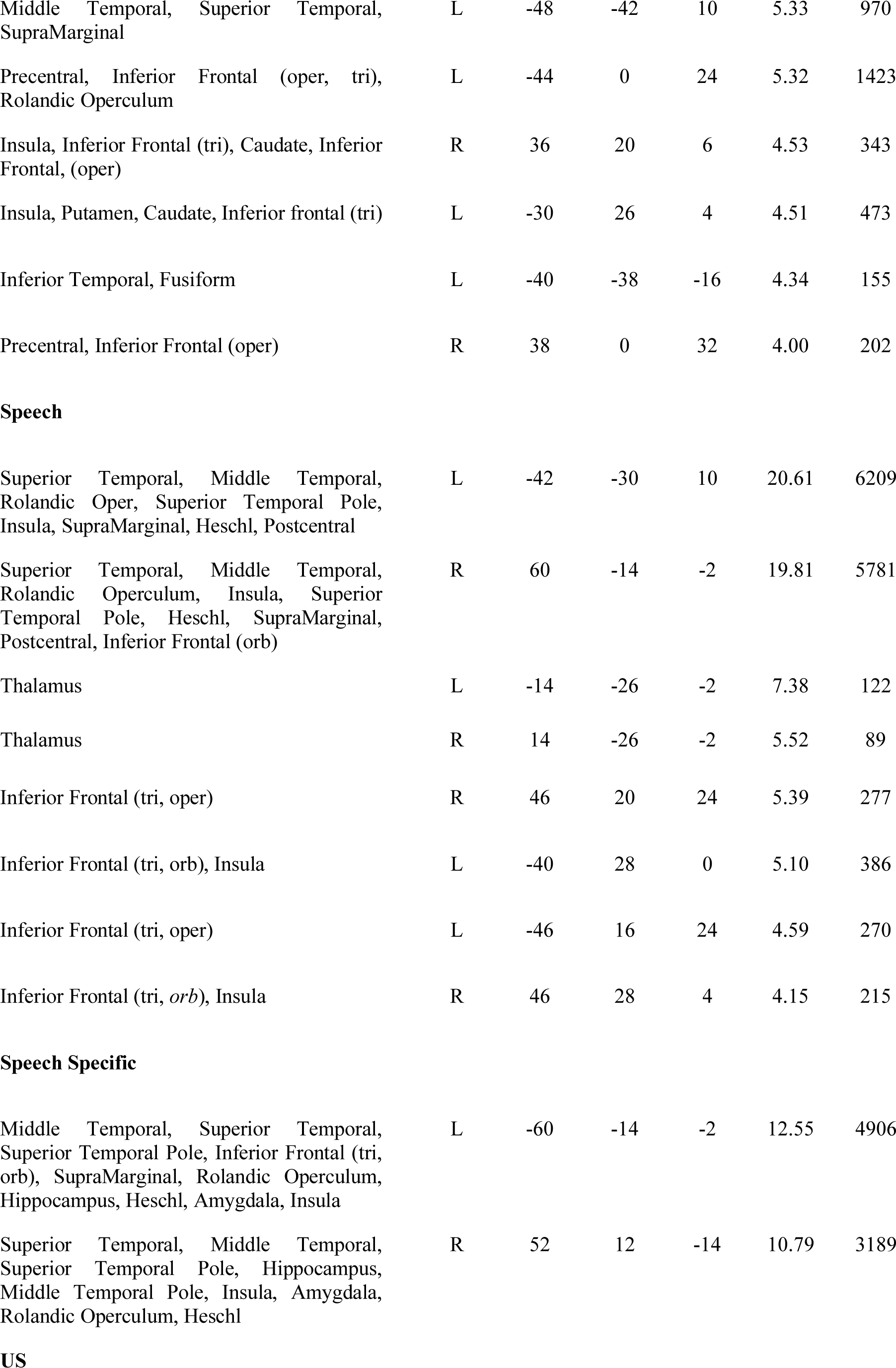
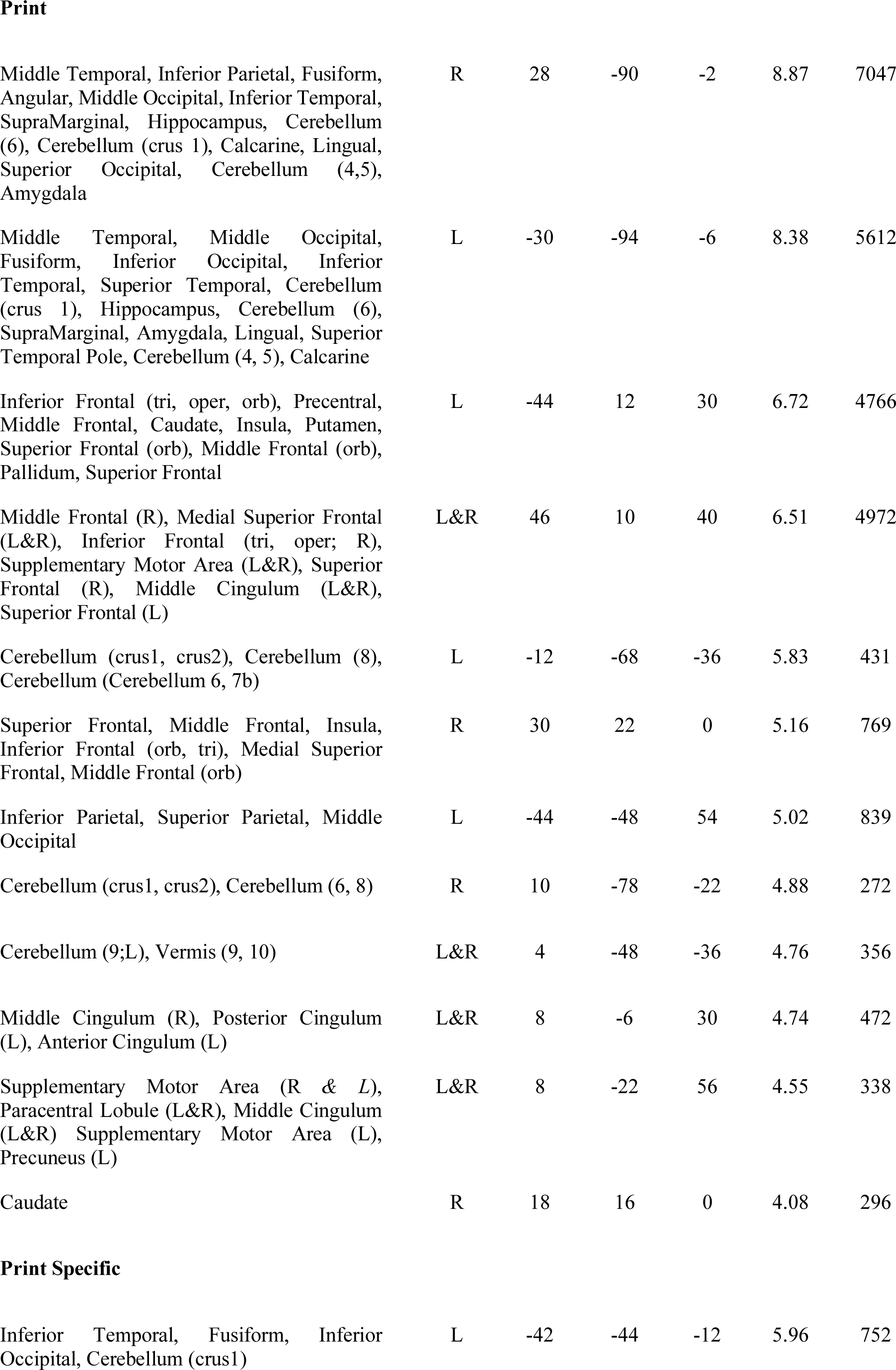
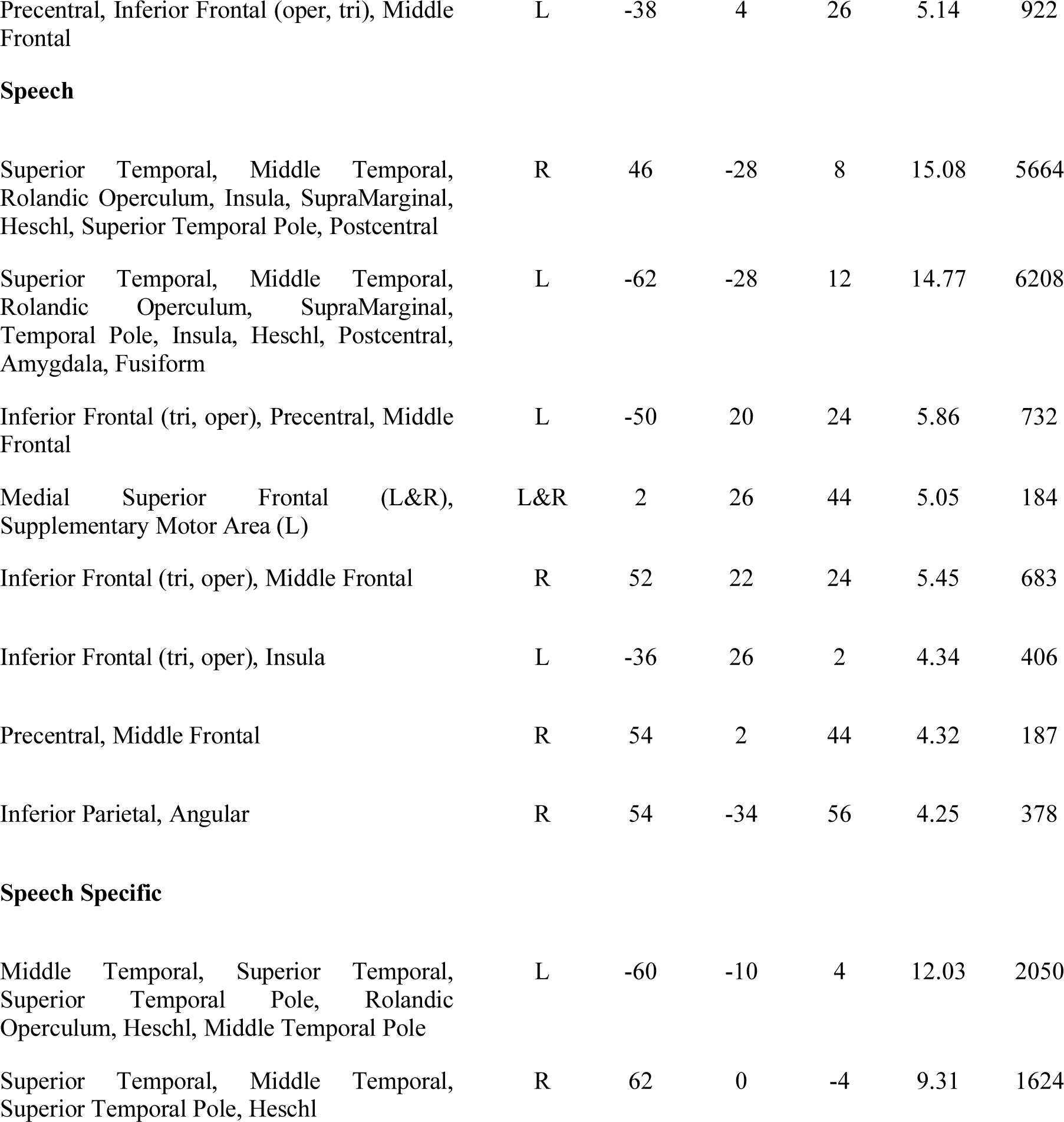
Brain regions that are active for Print, Speech, Print Specific (Print > Symbols) and Speech Specific (Speech > Vocoded). Threshold for each contrast p < 0.005, FDR-corrected. Hemisphere (H), coordinates (x, y, z), t-statistic for the peak (t) and number of voxels (v) are reported.

| Brain region | H | x | y | z | t | v |
| --- | --- | --- | --- | --- | --- | --- |
| **PL** |  |  |  |  |  |  |
| **Print** |  |  |  |  |  |  |
| Inferior Occipital, Middle Occipital, Inferior Temporal, Lingual, Middle Temporal, Cerebellum (crus 1), Cerebellum (6) | L | -38 | -64 | -12 | 11.42 | 2693 |
| Middle Temporal, Inferior Occipital, Superior Temporal, Fusiform, Inferior Temporal, Middle Occipital, Calcarine, Lingual, Cerebellum (4,5,6) | R | 30 | -90 | -6 | 10.55 | 3761 |
| Middle Temporal, Superior Temporal | L | -48 | -44 | 8 | 6.36 | 851 |
| Precentral, Inferior Frontal (oper, tri) | L | -42 | 10 | 30 | 5.91 | 1114 |
| Insula, Inferior Frontal (oper, tri), Precentral, Middle Frontal | R | 32 | 26 | 2 | 5.86 | 1243 |
| Supplementary Motor Area (L&R) | L&R | -10 | 8 | 50 | 5.81 | 356 |
| Superior Parietal, Inferior Parietal | L | -28 | -60 | 48 | 4.51 | 187 |
| **Print Specific** |  |  |  |  |  |  |
| Supplementary Motor Area (L&R), Middle Cingulum (L&R) | L&R | -8 | 8 | 50 | 6.70 | 838 |
| Superior Temporal, Middle Temporal, SupraMarginal | R | 44 | -40 | 18 | 5.38 | 989 |
| Middle Temporal, Superior Temporal, SupraMarginal | L | -48 | -42 | 10 | 5.33 | 970 |
| Precentral, Inferior Frontal (oper, tri), Rolandic Operculum | L | -44 | 0 | 24 | 5.32 | 1423 |
| Insula, Inferior Frontal (tri), Caudate, Inferior Frontal, (oper) | R | 36 | 20 | 6 | 4.53 | 343 |
| Insula, Putamen, Caudate, Inferior frontal (tri) | L | -30 | 26 | 4 | 4.51 | 473 |
| Inferior Temporal, Fusiform | L | -40 | -38 | -16 | 4.34 | 155 |
| Precentral, Inferior Frontal (oper) | R | 38 | 0 | 32 | 4.00 | 202 |
| Speech |  |  |  |  |  |  |
| Superior Temporal, Middle Temporal, Rolandic Oper, Superior Temporal Pole, Insula, SupraMarginal, Heschl, Postcentral | L | -42 | -30 | 10 | 20.61 | 6209 |
| Superior Temporal, Middle Temporal, Rolandic Operculum, Insula, Superior Temporal Pole, Heschl, SupraMarginal, Postcentral, Inferior Frontal (orb) | R | 60 | -14 | -2 | 19.81 | 5781 |
| Thalamus | L | -14 | -26 | -2 | 7.38 | 122 |
| Thalamus | R | 14 | -26 | -2 | 5.52 | 89 |
| Inferior Frontal (tri, oper) | R | 46 | 20 | 24 | 5.39 | 277 |
| Inferior Frontal (tri, orb), Insula | L | -40 | 28 | 0 | 5.10 | 386 |
| Inferior Frontal (tri, oper) | L | -46 | 16 | 24 | 4.59 | 270 |
| Inferior Frontal (tri, orb), Insula | R | 46 | 28 | 4 | 4.15 | 215 |
| Speech Specific |  |  |  |  |  |  |
| Middle Temporal, Superior Temporal, Superior Temporal Pole, Inferior Frontal (tri, orb), SupraMarginal, Rolandic Operculum, Hippocampus, Heschl, Amygdala, Insula | L | -60 | -14 | -2 | 12.55 | 4906 |
| Superior Temporal, Middle Temporal, Superior Temporal Pole, Hippocampus, Middle Temporal Pole, Insula, Amygdala, Rolandic Operculum, Heschl | R | 52 | 12 | -14 | 10.79 | 3189 |
| Print |  |  |  |  |  |  |
| Middle Temporal, Inferior Parietal, Fusiform, Angular, Middle Occipital, Inferior Temporal, SupraMarginal, Hippocampus, Cerebellum (6), Cerebellum (crus 1), Calcarine, Lingual, Superior Occipital, Cerebellum (4,5), Amygdala | R | 28 | -90 | -2 | 8.87 | 7047 |
| Middle Temporal, Middle Occipital, Fusiform, Inferior Occipital, Inferior Temporal, Superior Temporal, Cerebellum (crus 1), Hippocampus, Cerebellum (6), SupraMarginal, Amygdala, Lingual, Superior Temporal Pole, Cerebellum (4, 5), Calcarine | L | -30 | -94 | -6 | 8.38 | 5612 |
| Inferior Frontal (tri, oper, orb), Precentral, Middle Frontal, Caudate, Insula, Putamen, Superior Frontal (orb), Middle Frontal (orb), Pallidum, Superior Frontal | L | -44 | 12 | 30 | 6.72 | 4766 |
| Middle Frontal (R), Medial Superior Frontal (L&R), Inferior Frontal (tri, oper; R), Supplementary Motor Area (L&R), Superior Frontal (R), Middle Cingulum (L&R), Superior Frontal (L) | L&R | 46 | 10 | 40 | 6.51 | 4972 |
| Cerebellum (crus1, crus2), Cerebellum (8), Cerebellum (Cerebellum 6, 7b) | L | -12 | -68 | -36 | 5.83 | 431 |
| Superior Frontal, Middle Frontal, Insula, Inferior Frontal (orb, tri), Medial Superior Frontal, Middle Frontal (orb) | R | 30 | 22 | 0 | 5.16 | 769 |
| Inferior Parietal, Superior Parietal, Middle Occipital | L | -44 | -48 | 54 | 5.02 | 839 |
| Cerebellum (crus1, crus2), Cerebellum (6, 8) | R | 10 | -78 | -22 | 4.88 | 272 |
| Cerebellum (9;L), Vermis (9, 10) | L&R | 4 | -48 | -36 | 4.76 | 356 |
| Middle Cingulum (R), Posterior Cingulum (L), Anterior Cingulum (L) | L&R | 8 | -6 | 30 | 4.74 | 472 |
| Supplementary Motor Area (R & L), Paracentral Lobule (L&R), Middle Cingulum (L&R) Supplementary Motor Area (L), Precuneus (L) | L&R | 8 | -22 | 56 | 4.55 | 338 |
| Caudate | R | 18 | 16 | 0 | 4.08 | 296 |
| Print Specific |  |  |  |  |  |  |
| Inferior Temporal, Fusiform, Inferior Occipital, Cerebellum (crus1) | L | -42 | -44 | -12 | 5.96 | 752 |
| Precentral, Inferior Frontal (oper, tri), Middle Frontal | L | -38 | 4 | 26 | 5.14 | 922 |
| **Speech** |  |  |  |  |  |  |
| Superior Temporal, Middle Temporal, Rolandic Operculum, Insula, SupraMarginal, Heschl, Superior Temporal Pole, Postcentral | R | 46 | -28 | 8 | 15.08 | 5664 |
| Superior Temporal, Middle Temporal, Rolandic Operculum, SupraMarginal, Temporal Pole, Insula, Heschl, Postcentral, Amygdala, Fusiform | L | -62 | -28 | 12 | 14.77 | 6208 |
| Inferior Frontal (tri, oper), Precentral, Middle Frontal | L | -50 | 20 | 24 | 5.86 | 732 |
| Medial Superior Frontal (L&R), Supplementary Motor Area (L) | L&R | 2 | 26 | 44 | 5.05 | 184 |
| Inferior Frontal (tri, oper), Middle Frontal | R | 52 | 22 | 24 | 5.45 | 683 |
| Inferior Frontal (tri, oper), Insula | L | -36 | 26 | 2 | 4.34 | 406 |
| Precentral, Middle Frontal | R | 54 | 2 | 44 | 4.32 | 187 |
| Inferior Parietal, Angular | R | 54 | -34 | 56 | 4.25 | 378 |
| **Speech Specific** |  |  |  |  |  |  |
| Middle Temporal, Superior Temporal, Superior Temporal Pole, Rolandic Operculum, Heschl, Middle Temporal Pole | L | -60 | -10 | 4 | 12.03 | 2050 |
| Superior Temporal, Middle Temporal, Superior Temporal Pole, Heschl | R | 62 | 0 | -4 | 9.31 | 1624 |

**Table S2.**
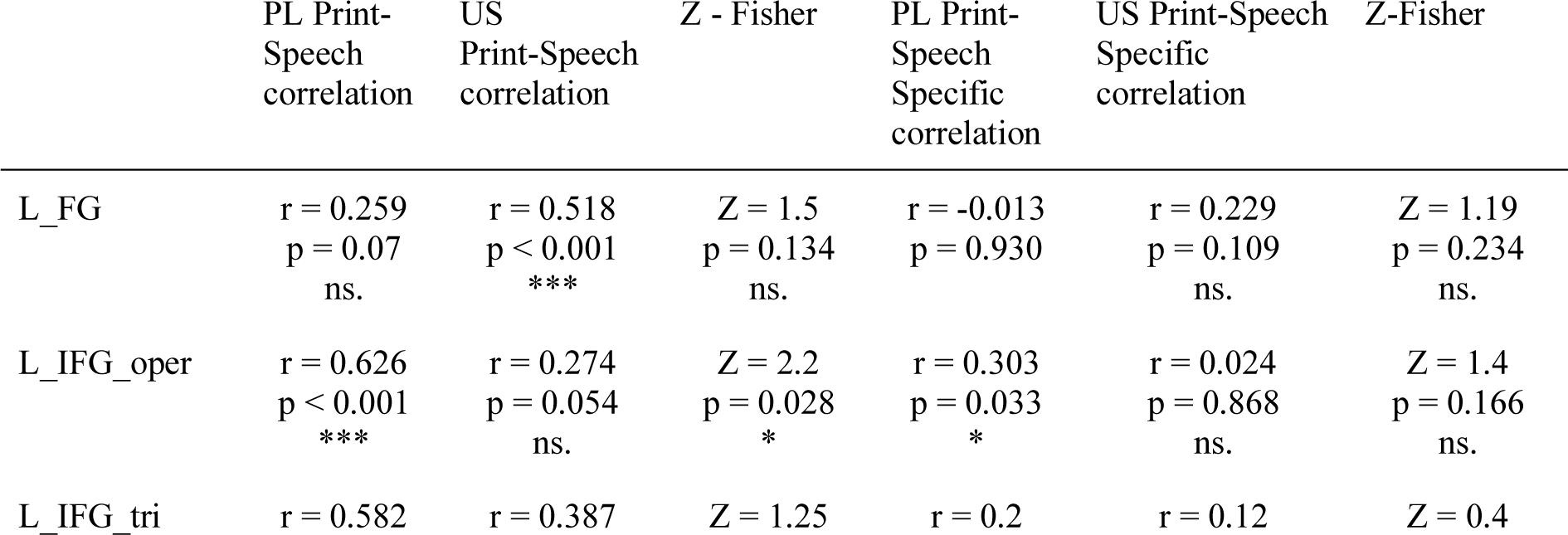
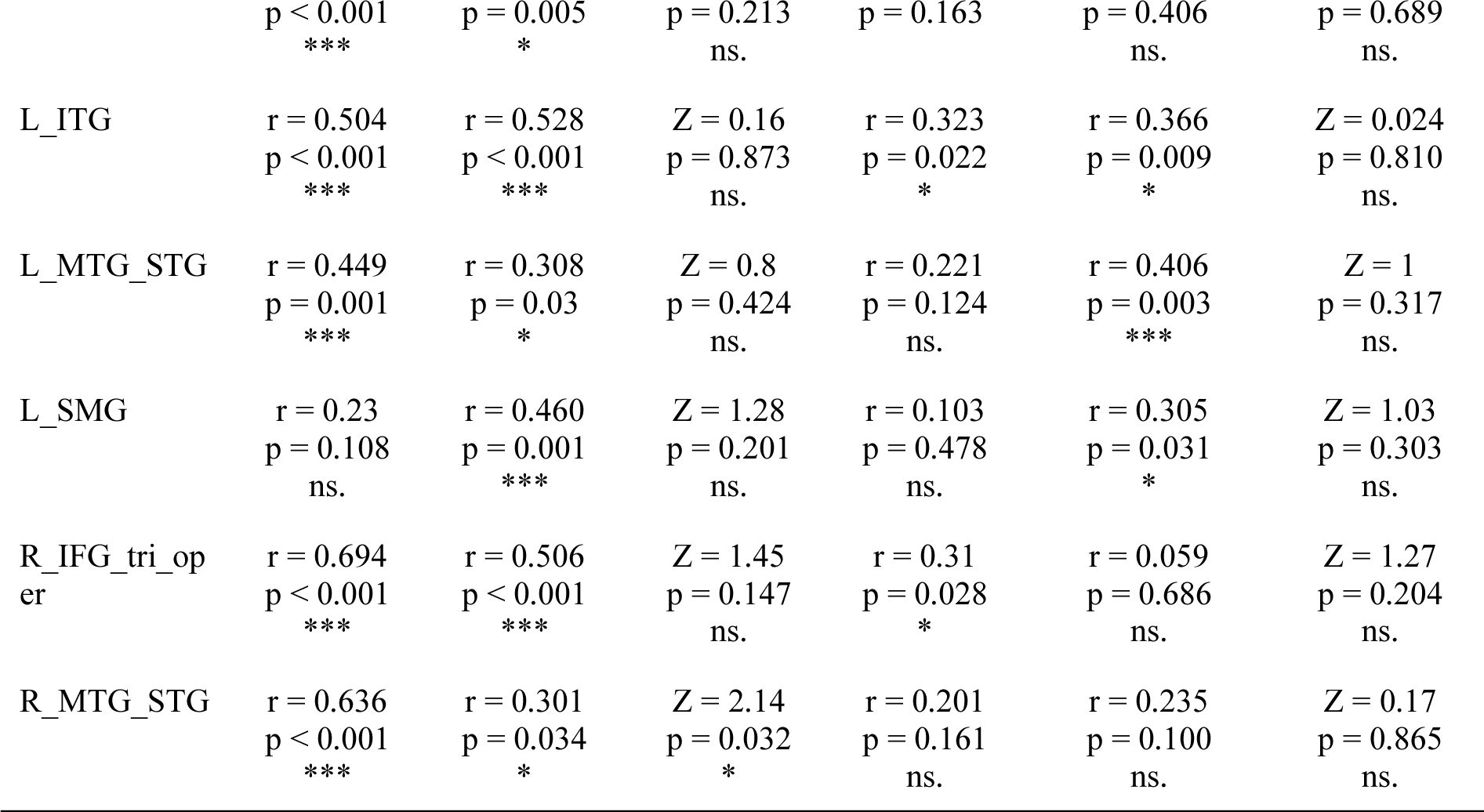
Print-Speech and Print-Speech Specific Correlations within ROIs.

